# The challenges of single cell transcriptomics on difficult human tissue: the placenta

**DOI:** 10.1101/2025.11.14.688483

**Authors:** Theodoros Xenakis, Jose J Moreno-Villena, George Hall, Sara L Hillman, Yara E Sanchez-Corrales, Sergi Castellano

## Abstract

Demand for the application of single cell transcriptomics on difficult tissues, processed and stored in disparate conditions, has led to the development of various single cell modalities. We focus on the placenta, a challenging tissue to transcriptomically interrogate due to senescence, intermittent hypoxia, high levels of RNase activity and tissue trauma at delivery. We performed single cell and nuclei transcriptomics on two samples with probe-based and native molecule transcript capture, droplet based and in situ plate based cell isolation. We explored sample and storage variations, including freshly dissociated tissues, fixed cells, snap frozen and FFPE processing. We find that variations in sample processing and storage have much larger effect on cell type proportions than differences in chemistry, impacting the biology that can be learned or the identification of disease markers. Further, the transcriptomic output of in situ combinatorial indexing consistently overlaps, with single nucleus transcriptomics, sharing little with other modalities. This may limit combinatorial indexing for sampling cytoplasmstic mRNAs. Our comprehensive analysis of the varying effect of single cell transcriptomic modalities, including their sample management strategies, provides novel and essential considerations for the experimental design and analysis of single cell transcriptomics applicable to challenging tissues, offering actionable guidance for future experiments.

**GRAPHICAL ABSTRACT:** 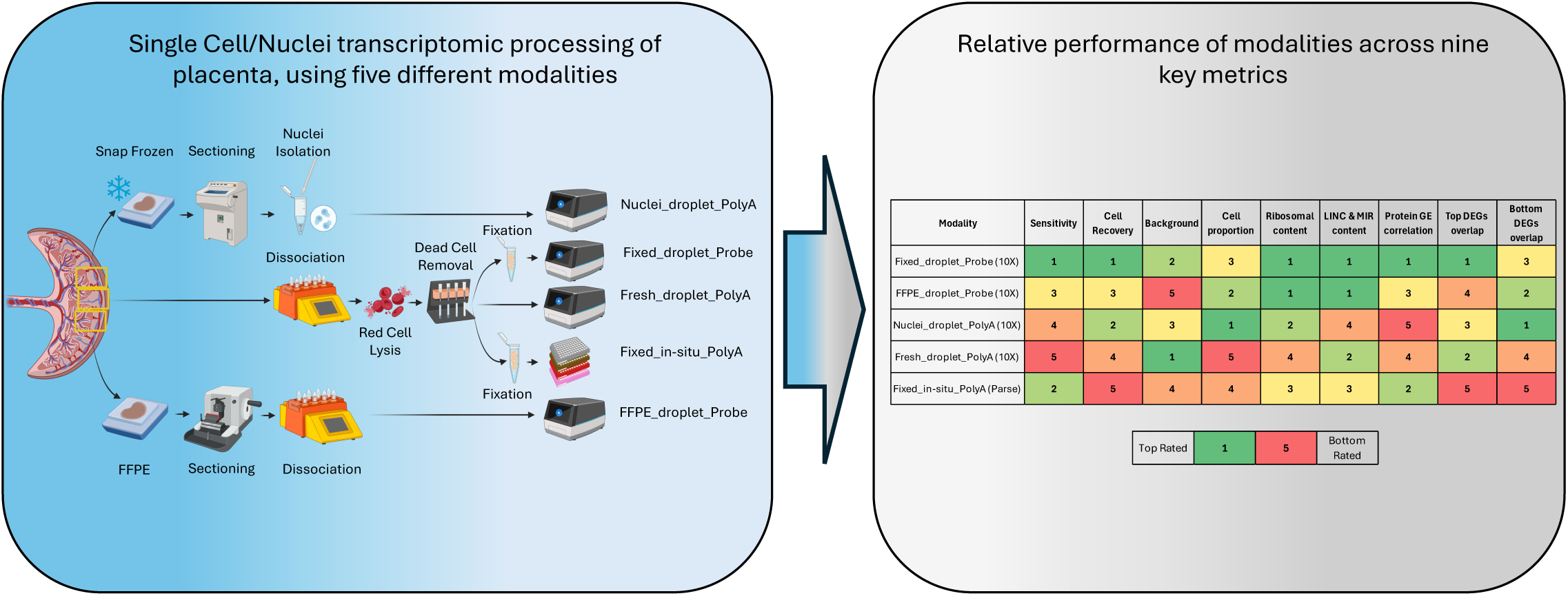

## INTRODUCTION

Biological tissues are inherently complex and heterogenous. They comprise of cell types and sub-types which can vary in their individual and functional states according to both external factors, such as cell singalling from other cells, and internal processes, such as cell cycle phase or apoptosis. The more so when tissues are affected by pathological changes [1]. Single cell transcriptomics quantifies this cellular heterogeneity, in health and disease, with promising applications in basic and translational research [2].

The popularity of single cell transcriptomics has justifiably increased, with rising demand for its application on increasingly difficult tissues with varying processing and storage conditions. Previously, freshly dissociated viable cells where the only option as an input in the vast majority of commercially available solutions, limiting the application of single cell transcriptomics. In recent years, methodological modalities have been developed to enable the single cell transcriptomic analysis of isolated nuclei from snap frozen tissue, cells fixed after tissue dissociation and cells isolated from formalin fixed paraffin embedded (FFPE) tissue blocks. This was greatly enabled by optimising transcript capture using probe sets that target protein-coding genes, as opposed to oligonucleotides that capture the 3’ prime poly-adenylated tails of mRNA molecules. Furthermore, novel plate-based methods promise the opportunity to circumvent cell-shape capture limitations inherent to droplet-based methods.

Previous comparative studies have mostly interrogated either naïve or sorted peripheral blood mononuclear cells (PBMCs) and cell lines. They have demonstrated the variance in performance and functional outputs using different commercially available kits, exploring their effects on sample multiplexing, case control comparisons and drug screening potential [3–10]. In an analogous fashion, commercially available solutions have been developed and optimised on similar common sample types such as PBMCs and then validated on other sample types, retrospectively.

Here, we focus on a more challenging tissue, the placenta, allowing us to better understand the limitations of different modalities in less optimal sample types. The placenta is the site of direct molecular and cellular communication between the pregnant woman and the fetus and, has traditionally been considered a difficult tissue to transcriptionally interrogate, particularly from pregnancies affected by pathology. This is due to senescence, intermittent hypoxia, high levels of RNase activity and tissue trauma at delivery [11, 12]. Understanding the effect of these issues can be more broadly applicable as similar difficulties have also been reported on pancreas and liver [13, 14]. These, and other tissues pose a challenge for transcriptomic solutions which rely on capturing native mRNA molecules from their 3’ prime poly-adenylated tail (poly-A). A challenge that may be overcome using near whole transcriptome probe sets.

Furthermore, early studies on single cell methodologies on the placenta have predominantly focused on differences between single nuclei and single cell methods, where important findings regarding cell type resolution and loss of disease signatures have been reported [15, 16]. However, appropriate representation of certain cell types in the placenta, such as the multinucleated syncitiotrophoblasts (STBs) [17], myeloid cell types remains challenging to achieve with droplet-based modalities due to cell shape and size restrictions [18]. These challenges have also affected the interrogation of binucleated cardiomyocytes in heart tissue and neurons in brain tissue, further highlighting the importance of understanding how different modalities can overcome these challenges [19, 20].

To understand the wider effect of different single-cell transcriptomic modalities on difficult tissues, including different sample storage and preprocessing, we performed single cell and nuclei transcriptomics on two samples using five different modalities. Interestingly, we report that sample storage and processing have significant impact on cell type population ratios. Furthermore, we present evidence that in situ combinatorial indexing consistently shares similarities in transcriptomic output with single nucleus transcriptomics and yet shows distinct lack of overlap with other modalities, of the most overexpressed genes. Though new versions of kits and updated chemistries may improve some of these limitations across modalities, our findings fundamentally relate to their underlying mechanisms, which in conjunction with the impact from processing and storing samples, are likely intrinsic to each single cell technology. Finally, we offer guidance to the design of single cell experiments for each modality in a simple way (**Figure 5b**).

## MATERIAL AND METHODS

### Patient recruitment

We recruited pregnant women at the fetal medicine unit, antenatal care unit and labour ward of the University College London Hospitals, NHS Foundation Trust. The study cohort comprised 3 pregnant women. The study was approved by the South Oxford Research Ethics Committee (REC CODE: 17/SC/0432).

Inclusion criteria were women with a singleton pregnancy, able to consent, experiencing preterm delivery by C-section due to iatrogenic reasons prior to 37 weeks of gestation. Exclusion criteria were women with pregnancies affected by major fetal anomalies (whether chromosomal, structural, or genetic), twin pregnancies and comorbidities including diabetes mellitus (gestational or T1 or T2), maternal autoimmune, renal or heart disease. Donors were excluded if had any clinical symptoms/signs of infection or a positive test of infection during standard clinical practice.

Placentas from two of the recruited patients were used for the single cell/nuclei modality comparison and the placenta of the remaining patient was used for in situ Spatial Transcriptomics (iST).

### Sampling

After delivery, the placentas were washed twice in 1x DPBS (-Ca, -Mg). The largest distance between the cord insertion site and the edge of the placenta was then established. From the mid-point of said distance three adjacent 2cm x 2cm samples were taken. One was placed in cold HypoThermosol FRS (Sigma-Aldrich, cat no: H4416) at 4°C for a maximum of 24 hours, for dissociation and single-cell sequencing. The second was snap frozen in an isopentane bath set in dry ice and embedded in OCT, and stored at -80°C. The third was placed in formalin saline solution for paraffin embedding to prepare FFPE blocks. All sampling occurred within 30 minutes of delivery to ensure RNA integrity.

### Dissociation

Samples were removed from HypoThermosol FRS and transferred to C-Tubes (Milteny Biotec, cat no: 130-093-237) on ice before being minced with scissors. 10ml of Accutase (Sigma-Aldrich, cat no: A6964) was then added to the C-Tubes. The tissues were then dissociated on a gentleMACS Octo dissociator using a custom protocol. Dissociated cell suspensions were then passed through stacked 100um and 70um cell strainers (Milteny Biotec, cat no: 130-098-463, 130-098-462), pre-wetted with 2ml of DMEM, 10% FBS. Cell strainers were then rinsed with a further 10ml of DMEM, 10% FBS.

Cells were pelleted by centrifugation at 300RCF for 5 minutes. The supernatant was then removed, and the pellets were resuspended in 20ml of Red Cell Lysis buffer (Myltenyi Biotec, cat no: 130-094-183) and incubated for 10 minutes at room temperature on a rocker. Then 30ml of cold 1x DPBS was added, and cells were pelleted by centrifugation at 300RCF for 5 minutes. The supernatant was removed, and cells were washed by resuspending in 5ml DMEM, 10% FBS before being pelleted by centrifugation at 300RCF for 5 minutes.

The supernatant was then removed, and cell pellets were resuspended in 1ml of Dead Cell Removal MicroBeads (Miltenyi Biotec, cat no:130-090-101) and incubated for 15 minutes at room temperature. Suspensions were then loaded onto MACS LS columns (Miltenyi Biotec, cat no:130-042-401) pre-rinsed with 3ml 1x Binding Buffer. Columns were rinsed with a further 3ml of Dead Cell Removal 1x Binding Buffer. Cells were then pelleted by centrifugation at 300RCF for 5 minutes. The supernatant was removed, and cells were resuspended in 1ml DMEM, 10% FBS.

Final cell suspensions for placental samples of both patients were divided into three. One to be used immediately with the 10X Genomics Next GEM Single-cell 5’ v2 kit (Fresh_droplet_Poly-A). The second was fixed according to the Parse Biosciences Evercode Cell Fixation v2 User Manual v2.3 to be used with the Parse Biosciences Evercode WT Mini v2 kit (Fixed_in-situ_Poly-A). The third was fixed according to 10X Genomics demonstrated protocol CG000478 rev D to be used with the 10X Genomics Chromium Fixed RNA Profiling Reagent Kit (Fixed_droplet_Probe).

### Nuclei Isolations from placenta samples

Snap frozen, OCT embedded placental samples were cryosectioned on a Bright OTF5000 cryostat. Five 25um sections were collected. Nuclei were isolated from the sections, using Chromium Nuclei Isolation Kit (10X Genomics, cat no: 1000494), following 10X Genomics user guide CG000505, to be used with 10X Genomics Next GEM Single-cell 5’ v2 kit (Nuclei_droplet_Poly-A).

### FFPE dissociations from placental samples

FFPE blocks from placenta samples of both patients were sectioned on a microtome. Five 25um sections were collected and dissociated according to 10X Genomics demonstrated protocol CG000632 revD to be used with the 10X Genomics Chromium Fixed RNA Profiling Reagent Kit (FFPE_droplet_Probe).

### Generating single cell transcriptomic libraries

Dissociated cells and isolated nuclei were checked for concentration and viability using an Acridine Orange/Propidium Iodide Stain (logos Biosystems, cat no: F23001) on a logos Biosystems Luna FL automated cell counter. Cells/nuclei for each sample were used to generate single cell/nuclei transcriptomic libraries according to the specifications listed in Table 1.

**Table 1.** Library preparation details per single cell modality.

| Modality | Cell origin | Single-cell sequencing kit | Library Prep User Guide followed |
| --- | --- | --- | --- |
| **Fresh_droplet_PolyA** | Cells dissociated from fresh tissue | Chromium Next GEM Single-cell 5' Kit v2 kit | 10X Genomics CG000331 |
| **Nuclei_droplet_PolyA** | Nuclei from snap frozen tissue | Chromium Next GEM Single-cell 5' Kit v2 kit | 10X Genomics CG000331 |
| **FFPE_droplet_Probe** | Cells dissociated from FFPE blocks | Chromium Fixed RNA Profiling Reagent Kit | 10X Genomics CG000691 |
| **Fixed_droplet_Probe** | Cells dissociated from fresh tissue then fixed | Chromium Fixed RNA Profiling Reagent Kit | 10X Genomics CG000691 |
| **Fixed_in-situ_PolyA** | Cells dissociated from fresh tissue then fixed | Evercode WT Mini v2 kit | Parse Biosciences Evercode WT Mini v2 User Manual v2.3. |

### Sequencing

Resulting single cell and single nuclei transcriptomic libraries were sequenced on an Illumina Novaseq 6000 sequencer using S4 (200 cycle) v1.5 sequencing kits (Illumina, cat no: 20028313). Libraries were sequenced over the minimum recommended depth per library type.

### In situ Spatial Transcriptomics

An 8um section of placenta was loaded onto specialised slide to be processed on the 10X Genomics Xenium Analyser (10X Genomics, cat no’s: 1000460, 1000487, 1000461) using the Xenium Human Multi-Tissue and Cancer Panel (10X Genomics, cat no: 1000626) and a Xenium Add-on Custom 51 to 100 Gene Panel (10X Genomics, cat no: 1000651) following 10X Genomics demonstrated protocol CG000581 and user guides CG000582 and CG000584. The gene selection for the Add-on Custom 51 to 100 Gene Panel was based on differentially expressed genes in STB and CTB from a previous study [21].

### Count matrix generation and background removal

Droplet based: For each single cell and single nucleus library, the raw bcl files were converted into fastq files using cellranger “mkfastq”. Reads were aligned to the GRCh38-2020-A human reference genome (GENCODE v32/Ensembl98 distributed by 10X Genomics) and a matrix of unique transcripts per cell for each library was obtained using cellranger “multi”. We used CellBender 0.2.0 [22] to correct for ambient RNA per library and classify cell-containing droplets from empty ones.

In situ: For each single-cell library, the raw bcl files were converted into fastq files using bcl2fatsq. To minimise bias in gene detection and ensure the fidelity of the comparisons across all single-cell transcriptomic modalities, reads were aligned again to the GRCh38-2020-A human reference genome (GENCODE v32/Ensembl98 distributed by 10X Genomics) and a matrix of unique transcripts per cell for each library was obtained using split-pipe “all” and “comb”. In order to appropriately calculate ambient RNA, using Seurat 5.1.0 [23] we merged the unfiltered library matrixes and generated a .h5ad object. We then used CellBender 0.2.0 to correct for ambient RNA.

Following matrix generation and background removal the libraries of each modality were integrated, filtered for low quality cells and subjected to cell type annotation separately with the exception of FFPE_droplet_Probe and Fixed_droplet_Probe which were processed together.

### Quality Control, Integration and cell type annotations

We filtered low quality cells using Seurat 5.1.0. We excluded cells with number of detected genes below 400 and above 6,000, with transcript counts above 17,500, with percentage of mitochondrial reads more than 10% and percentage of haemoglobin genes more than 0.25..

Pre-processed data files generated from the Xenium instrument on-board analysis software were used to generate Seurat objects using Seurat, filtering out cells with 0 transcript counts.

Data integrations were performed using a standard workflow, including normalisation (“LogNormalise”), scale (“ScaleData”) for single-cell/nuclei libraries and (“SCTransform”) for iST. Highly variable genes selection was used to perform principal component analysis, k-nearest-neighbour calculation and graph-based community detection using Louvain clustering

We annotated cell types using gene expression of manually curated genes from the literature. Cell types were annotated at two levels of resolution. First, simply into immune and non-immune groups. Subsetting each group and repeating the clustering pipeline allowed for another level of annotation with higher granularity. During clustering, we detected doublet clusters (identified by a mixed profile of mutually exclusive marker genes for different cell types) and those were also excluded.

### Cell type proportions, gene expression metrics and cell capture rates

Filtered and annotated libraries were then merged using Seurat 5.1.0, and cell type annotations were checked for consistency. Cell type proportions for each method where then calculated for both levels of annotations. Percentage of gene expression per cell for three different classes of genes were calculated for each modality. These were Long Intergenic Non-Coding (LINC), Ribosomal Protein (RP) and Micro RNA (MIR) genes. The number of cells remaining post quality and doublet filtration was then calculated as a percentage of the number of cells inputted during sample processing.

### Principal component analysis

The most consistently represented cell type across all modalities were the STB cells. As such using Seurat, we subsetted the STB cells, and then pseudobulked (“AggregateExpression”) their gene expression by sample. We then ran Principal Component Analysis (“RunPCA”) using the top 2,000 most variable genes per library with npcs=8 and visualised the top 5 principal components in a UMAP plot (“RunUMAP”).

### Pseudo-bulking and gene expression correlation

The most represented cell type amongst the immune cell populations across the five modalities were the NK/T cells. Conversely from the non-immune population, as mentioned previously, the most represented cell type were the STB cells. Using Seurat, we subsetted these cell types, and gene expression matrixes were subsetted to only include the genes that were shared between the probe set and poly-A capture approaches. We then pseudobulked (“AggregateExpression”), per cell type and modality, generating a per cell type gene expression matrix for each modality. A correlation matrix was then calculated from the normalised transcript counts.

### Inter-method differential gene expression

To assess the overlap of differential gene expression (DEG) analysis outputs we again subsetted the STB cells. We performed pairwise Wilcoxon rank-sum test analysis (“FindMarkers”) between the two biological replicates within each modalityretaining only DEGs with an adjusted p value below 0.05 and ranked them by descending log2 fold change. We subsequently calculated the ratio of overlap for the top (most overexpressed) 10, 25, 50 and 100 DEGs independently as well as the bottom (most underexpressed) 10, 25, 50 and 100 DEGs.

## RESULTS

### Assessment of single cell transcriptomic modalities

To examine the varying effect of five different single cell modalities on the transcriptome, we processed two placental samples, which included the maternal decidua (**Figures 1a** and **b**). These two samples where from late preterm deliveries, delivered at 37 and 36 gestational weeks respectively. We appreciate that the low sample number in this study is a limitation, however, we consistently report low variance between the two samples in our analyses (**Figures 2a, 4a** and **b, Extended Figures 1 and 3**).

**Figure 1:**
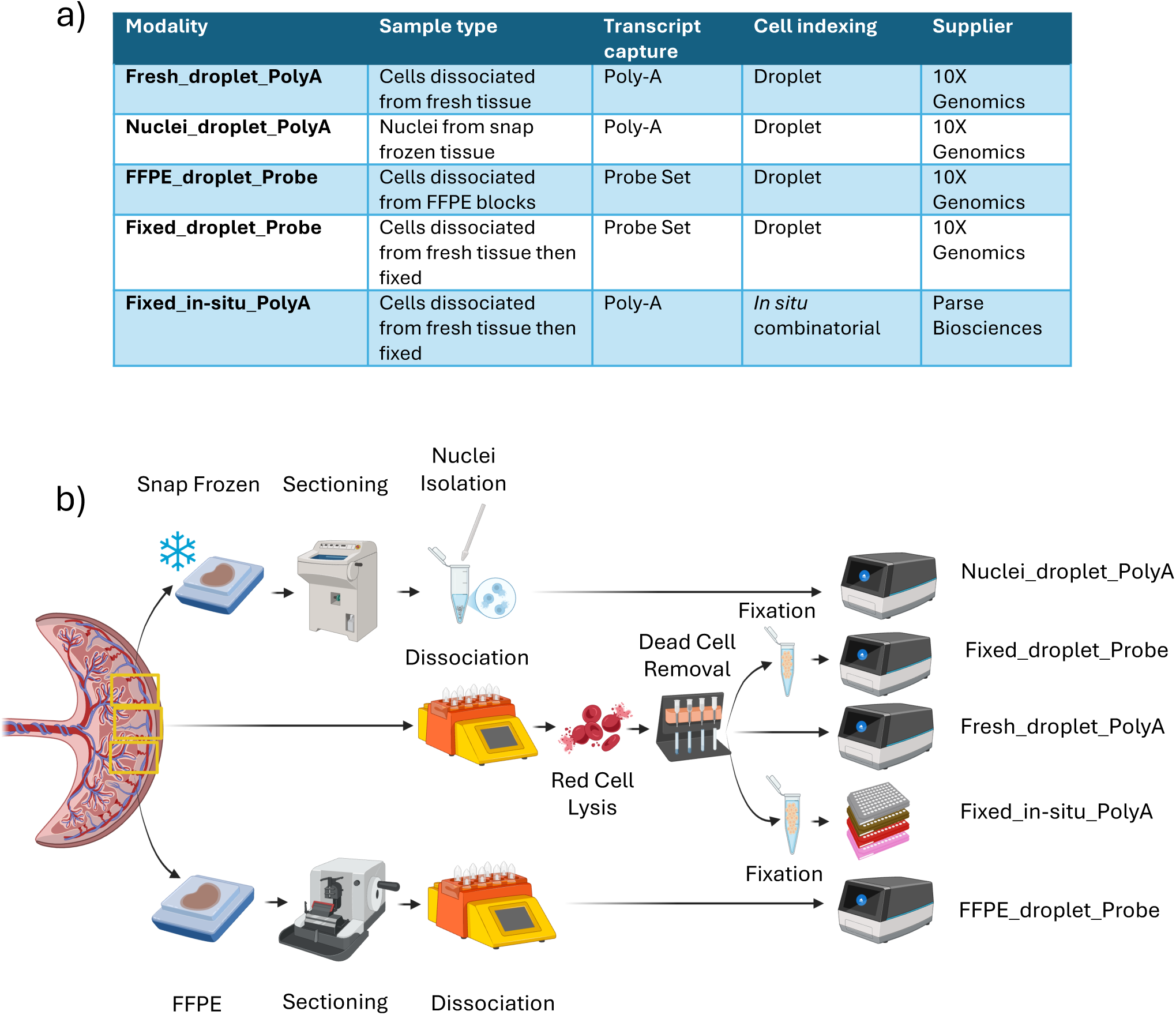
Overview of single cell/ nuclei modalities. **a**) Table detailing the variable elements of the five different modalities interrogated. **b**) Graphical representation of sampling and key processing steps for each modality.

**Figure 2:**
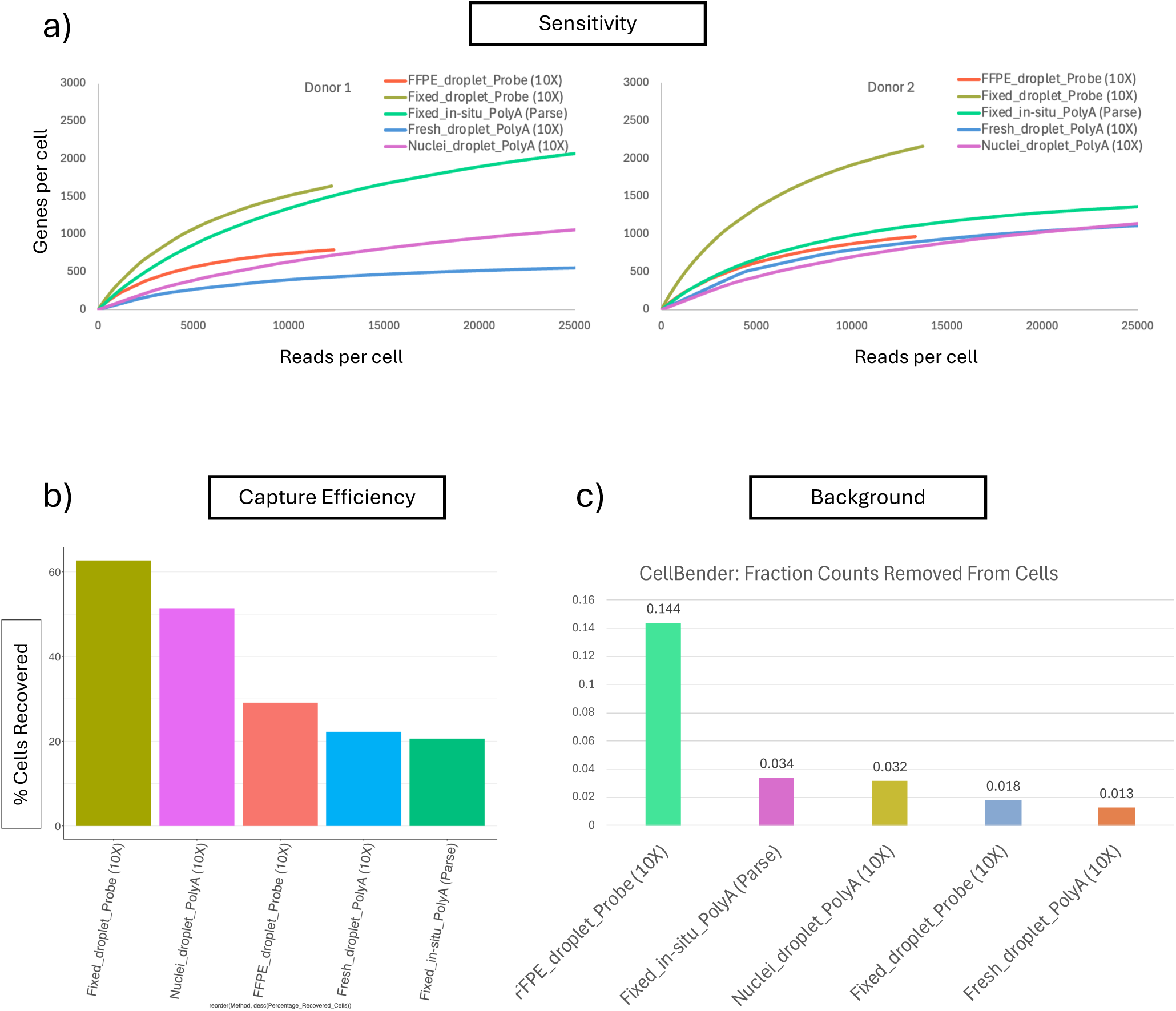
Performance metrics of modalities. **a**) Sensitivity, as illustrated for each donor using the relationship between median genes per cell and median reads per cell for the five modalities (sequencing depth shown for each modality is to the manufacturers minimum recommendation + 5%). **b**) Capture efficiency, showing the percentage of cells used for analysis post Quality Control filtering in relation to cells inputted for each modality. **c**) Background, illustrating per modality the fraction of reads removed by the computational tool used for removing reads assigned to ambient RNA.

**Figure 3:**
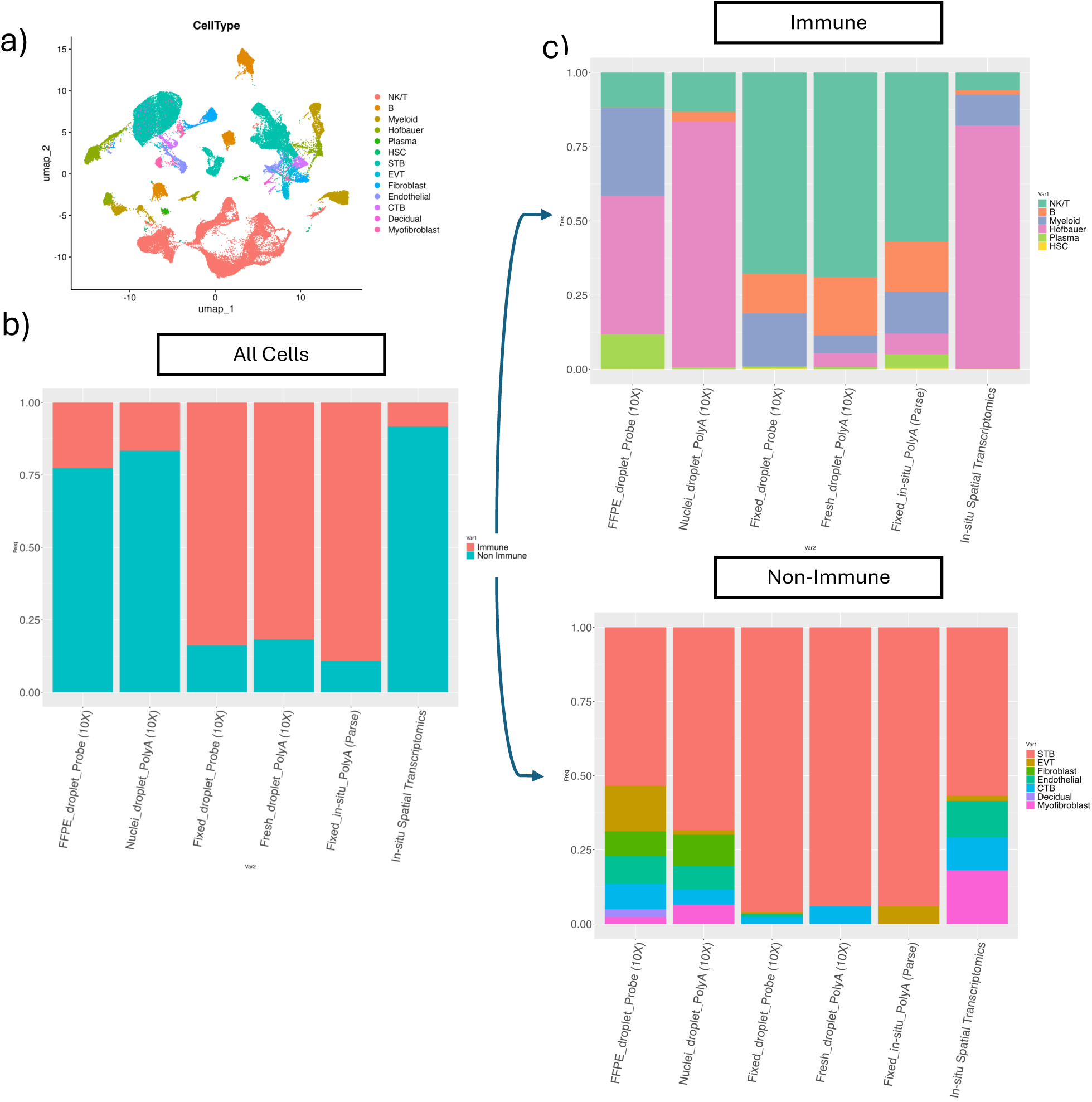
Cell type proportions. **a**) UMAP of combined dataset including all five modalities coloured by annotated cell type. **b**) Relative proportion of cells annotated as immune and non-immune cells by modality. **c**) Relative proportions of cell annotated within the immune and non-immune cell types.

**Figure 4:**
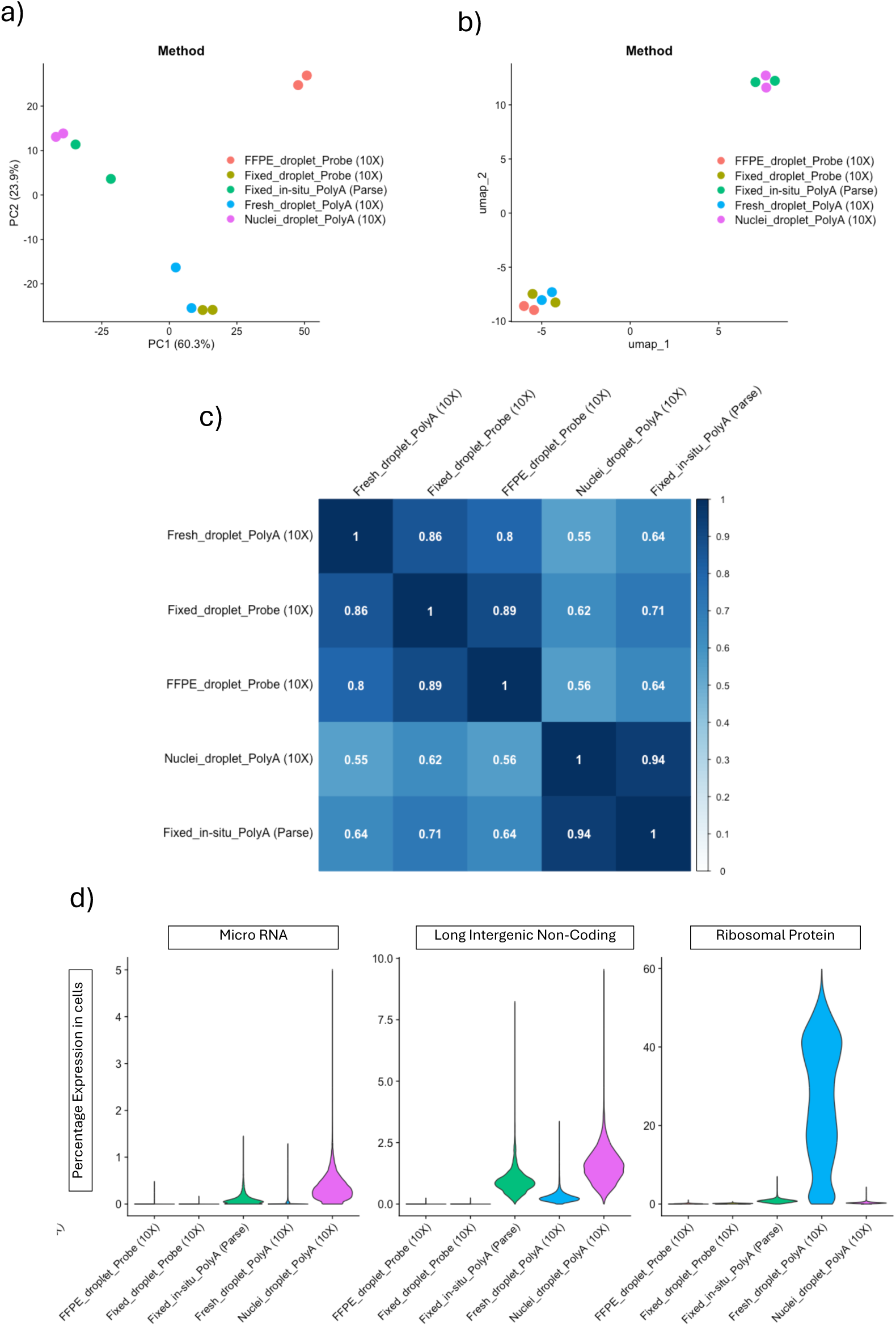
Gene expression differences across modalities. **a**) Principal Component Analysis of pseudobulked Syncitiotrophoblasts (STBs) expression of protein coding genes, per library, coloured by modality. **b**) UMAP (using top 5 Principal Components) of pseudobulked STB expression of protein coding genes, per library, coloured by modality. **c**) Correlation matrix of per cell type pseudobulked protein coding gene expression of STB and Natural Killer/T-cell across modalities. coloured by modality. **d**) Distribution of percentage expression in cells of Long Intergenic Non-Coding, microRNA and Ribosomal Protein genes per modality.

### Chemistry specific metrics

In the first instance, we examined the sensitivity of each modality by looking at the genes per cell in relation to the sequencing reads per cell (**Figure 2a**). In both samples, the highest sensitivity was observed in Fixed_droplet_Probe, perhaps unsurprisingly given the probe-based transcript capture. The second most sensitive modality is Fixed_in-situ_PolyA, meaning the top two modalities utilise cells fixed after dissociation of fresh tissue.

To understand the cell capture efficiency of each modality, we quantified the percentage of cells remaining (from the total number of cells used as an input) after ambient RNA removal (background) and quality filtering (**Figure 2b**). The highest cell capture efficiency was observed again in Fixed_droplet_Probe (60%) followed by Nuclei_droplet_PolyA (47%), where the capture efficiency is nearly triple and double, respectively, of the remaining three modalities. To understand the level of transcriptomic background introduced, we extracted the “fraction counts removed from cells” metric outputted from the computational background removal tool used (**Figure 1c**). The highest background was observed in FFPE_droplet_Probe (0.144%). This is in line with our expectations due to the challenging dissociation protocol required and the degraded nature of RNA from cells in FFPE tissue blocks. We observed good concordance between samples in both cell capture efficiency and background (**Extended Figure 1**). These results offer important information for the consideration of experimental designs.

### Proportions of cell types

After annotating the cell types represented in our final dataset (**Figure 3a**, **Extended Figure 2a**), we explored the proportion of each cell type per modality. We included carefully annotated in situ spatial transcriptomics (iST) from a placental sample of similar gestational age (35 weeks), in order to compare against a sample whose in vivo cell type proportions have been maximally maintained, due to the maintenance of tissue architecture iST (**Extended Figure 2b** and **c**). Initially, we annotated cells into immune and non-immune cell types (**Figure 3b, Extended Figure 3a**). Interestingly, the three modalities with cells from freshly dissociated tissue (Fresh_droplet_PolyA, Fixed_droplet_Probe and Fixed_in-situ_PolyA) consistently favoured immune cell types. In contrast, the two modalities with cells or nuclei from stored tissue blocks (FFPE_droplet_Probe and Nuclei_droplet_PolyA) consistently favoured non-immune cell types, which is in agreement with the iST sample.

We then investigated the immune and non-immune populations individually (**Figure 3c, Extended Figure 3b** and **c**), to explore cell types at higher granularity. In the immune population, again the three modalities with cells from freshly dissociated tissue (Fresh_droplet_PolyA, Fixed_droplet_Probe and Fixed_in-situ_PolyA) were relatively consistent for a lymphoid derived cell type (NK/T cells), which represented over half the cells in the population. This cell type is less present in the remaining two modalities (FFPE_droplet_Probe and Nuclei_droplet_PolyA), which is again consistent with the iST sample. FFPE_droplet_Probe has a broader representation of cell types, while Nuclei_droplet_PolyA greatly favours a fetal myeloid cell type (Hofbauer cells).

In the non-immune cell populations, there is again consistency in the three modalities with cells from freshly dissociated tissue (Fresh_droplet_PolyA, Fixed_droplet_Probe and Fixed_in-situ_PolyA), in which over 90% of cells in this population are syncitiotrophoblasts (STB) in each modality. There is, however, variation in the cell types in the remaining 10% of the non-immune population. Of note is the lack of myofibroblasts in these modalities, a cell type well represented in the iST sample. FFPE_droplet_Probe and Nuclei_droplet_PolyA both also have high representation of STBs; however, both have much broader representation of cell types, consistent with the iST sample.

These results showed good consistency between samples and underscore the need to understand the limitations of different methods of tissue processing.

### Protein coding gene expression variance

To assess how different modalities varied trascriptionally, we first applied dimensionality reduction techniques. We selected the cell type that is most consistently represented across all libraries (STB). To maintain consistency with most common analytical approaches we independently pseudo-bulked the expression profiles per library and subsetted the non-probe-based modalities to only include the genes that were present in the probe sets. This ensured we were looking at protein coding genes, avoiding biases caused by differences in gene sets explored. We then performed a principal component analysis, using the top 2,000 most variable genes per library (**Figure 4a**). The first principal component (PC1) accounted for 60.3% of the variance between libraries with the Nuclei_droplet_PolyA modality showing the largest amount of separation from the other modalities, followed by the Fixed_in-situ_PolyA. The second principal component (PC2) accounted for 23.9% of the variance with FFPE_droplet_Probe showing the largest amount of separation. The smallest separation between modalities is between Fixed_droplet_Probe and Fresh_droplet_PolyA. Intriguingly, despite the low variance between samples in each modality, the two libraries from Fixed_in-situ_PolyA and Fresh_droplet_PolyA show considerably higher separation across the two principal components compared to the separation between libraries produced by any other modality. This is echoed in the larger variance in sensitivity between biological replicates seen in Fixed_in-situ_PolyA and Fresh_droplet_PolyA which may indicate variation in output consistency between modalities. We then generated a UMAP, using the top 5 principal components (**Figure 4b**), in which we observed the Nuclei_droplet_PolyA clustering with the Fixed_in-situ_PolyA with clear separation from a cluster containing the remaining three modalities.

Furthermore, to quantifing the similarity of cell type-specific gene expression profiles among modalities, we selected the two cell types that are most consistently represented across modalities (STB and NK/T cells). We independently pseudo-bulked their expression profiles per modality for each cell type. We again subsetted the non-probe-based modalities to only include the genes that were present in the probe sets, and therefore protein coding. We then generated a correlation matrix from the normalised counts across modalities (**Figure 4c, Extended Figure 5a** and **b**). This allows us to assess the similarity of gene expression profiles from the same cell types across modalities using the same gene set. As expected, there was high correlation (Pearson coefficients: 0.8-0.89) between the three whole cell droplet-based modalities (Fresh_droplet_PolyA, Fixed_droplet_Probe and FFPE_droplet_Probe). The highest correlation (Pearson coefficient: 0.94), however, was between Fixed_in-situ_PolyA and Nuclei_droplet_PolyA. This echoes the PCA and UMAP results detailed above and indicates potential bias towards nuclear transcripts for the Fixed_in-situ_PolyA.

### Alternative gene expression features by modality

We also sought to understand the variance in the alternative genes detected between the different modalities. We queried the percentage of counts in cells per modality for three different classes of genes, that are not traditionally investigated in most transcriptomic analyses: ribosomal protein genes (RP), microRNA genes (MIR) and long intergenic non-coding genes (LINC) (**Figure 4d**). As MIR and LINC genes are non-coding genes and RP genes are highly conserved with low variance in expression, they are frequently excluded from analysis. By quantifying their prevalence in our results, we can assess the proportions of transcripts that are likely to be commonly excluded from investigations. Alternatively, where LINC and MIR genes are of particular interests, our quantification provides information on which modalities provide best coverage of these gene types. Fresh_droplet_PolyA had very high proportion of RP genes compared to the rest of the modalities with a significant number of cells reaching and even exceeding 40% of counts. The highest percentage of MIR and LINC genes was observed in Nuclei_droplet_PolyA, which is expected as both these gene classes are of nuclear RNAs. However, significant levels were also detected in Fixed_in-situ_PolyA. Unsurprisingly, both probe-based approaches (Fixed_droplet_Probe and FFPE_droplet_Probe) had almost no counts for any of the three gene classes, as probes targeting these genes are largely absent from their probe sets.

### Commonalities of differential gene expression analysis

To explore how different single cell modalities impact the results of differential gene expression analysis, we compared the same cell type (STBs) between two donors (biological replicates) in each modality. We filtered the resulting differentially expressed genes (DEGs), retaining only those with an adjusted p-value below 0.05 and ranked them by descending log2 fold change (**Extended Figure 4**). We subsequently explored the percentage of overlapping DEGs when looking at the top (most upregulated) and bottom (most downregulated) 10, 25, 50 and 100 DEGs (**Figure 5a**). We show that when exploring the top 10, Fixed_in-situ_PolyA has considerably lower overlapping DEGs across modalities. Athough this pattern does persist, as we look at increasing numbers of DEGs, the overlap does increase. Additionally, there is consistently high overlap between the two probe-based approaches (FFPE_droplet_Probe and Fixed_in-situ_PolyA) in both top and bottom DEGs. For the top 50 and 100 DEGs there is notable overlap between Fixed_droplet_Probe and Fresh_droplet_PolyA, supporting our findings in the PCA (**Figure 4a**). At the top 10 and 25 DEGs we also observe notable overlap between Fresh_droplet_PolyA and Nuclei_droplet_PolyA. Finally, when exploring the bottom 25, 50 and 100 DEGs there is consistently higher overlap between Nuclei_droplet_PolyA and Fixed_in-situ_PolyA.

**Figure 5:**
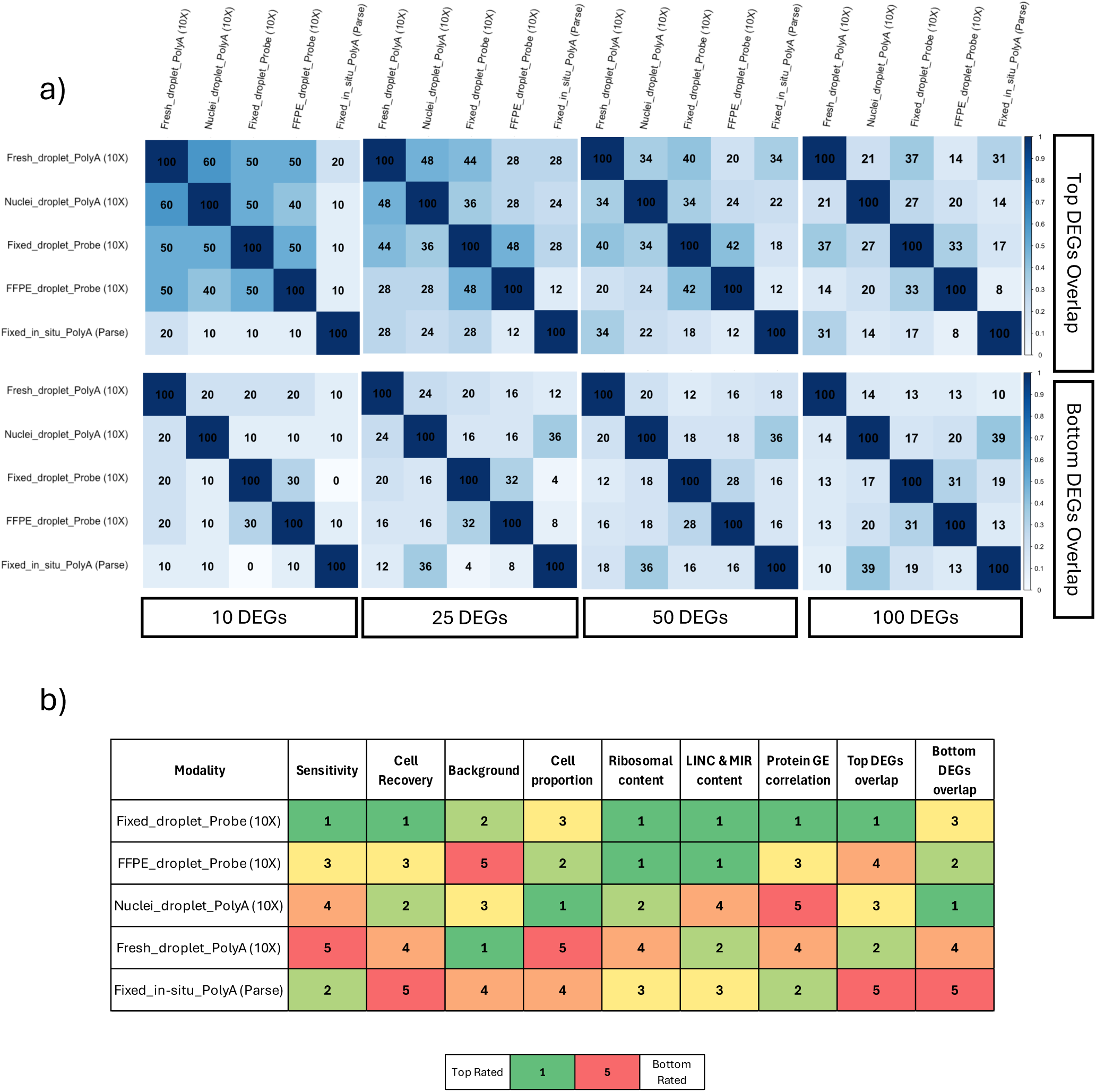
Differential gene expression analysis variance. **a**) Matrixes illustrating proportional overlap of 10, 25, 50 and 100 most over and under expressed genes by average log2 fold change (adjusted p-value <0.05), when comparing Syncitiotrophoblasts (STBs) between biological replicates, within each modality. **b)** Ranking of each modality by its performance on sensitivity, cell recovery, background amount, cell proportion (as judged by similarity to in-situ Spatial Transcriptomics), ribosomal protein content, LINC and MIR gene content, protein coding gene expression (GE) correlation (judged by average correlation to other modalities) and overlap of top and bottom Differentially Expressed Genes (DEGs) (judged by average overlap at 10, 25, 50 and 100 genes with other modalities).

### Overview of different modalities

In order to provide a summary of key metrics that are of significant importance for experimental design and analytical approaches, we present an overview of the comparable performance of the modalities explored across nine variables (**Figure 5b**). This can be used as a simple guide to the design of future experiments according to sample, storage and processing characteristics.

## DISCUSSION

We present a comparison of five single cell and nuclei transcriptomic modalities on a challenging tissue, the placenta. Though all modalities yielded reasonable results, for both biological replicates, the highest sensitivity is achieved with Fixed_droplet_Probe. Due to the probe-based transcript capture excluding non-protein coding genes, as well as other gene biotypes, the supplier’s recommended sequencing depth is also half of that of the polyA modalities, making it a more cost-effective solution. An important consideration, however, is the absence of some genes from these probe sets. Probes that target genes with hypervariable regions such as human leukocyte antigen (HLA*), T-Cell receptor variable (TRV*) and immunoglobulin variable (IG*) genes are absent. For some biological questions this can be a serious limitation, as for example the expression of HLA-G, is a marker gene for extravillous trophoblasts (EVTs), is not interrogated by these probe sets. Custom probes for specific absent genes of interest can be used, but as with any custom addition, may yield suboptimal results. For exploratory investigations and for questions with immunological nuance, this poses a challenge.

A key motivation for utilising single cell transcriptomics is to quantify cell type proportions and resolve heterogenous gene expression among cell types. Thus, it is important to evaluate the capacity and potential bias of each modality to transcriptionally represent the different cell types. Our results delineate differences among modalities in cell type proportions. Notably, we identify tissue processing as the main driver of differing cell type proportions across modalities. The three modalities with cells from freshly dissociated tissue have broadly comparable cell type proportions, both at the lower and higher levels of granularity of cell type annotation. These are distinct from the two modalities with cells and nuclei from stored tissue blocks, either fresh frozen or FFPE, as well as the iST sample with relatively well-maintained in vivo cell type proportionality. This is likely due to the requirement for cells from freshly dissociated tissues to be highly viable (optimally over 90%) and free from red cell contamination. As such, following an enzymatic and mechanical dissociation, these cell suspensions are subjected to red cell lysis and Magnetic-Activated Cell Sorting (MACS) for dead cell removal. These processing steps undoubtedly affect cell type proportions, with more sensitive cell types adversely affected by red cell lysis and more resilient ones favourably enriched by the dead cell removal. Previous studies have reported different cell type proportions from single cell analyses from freshly dissociated placentas, using alternative dissociation protocols and samples with different biological variables. This further supports our claim that sample processing should be central to considerations of experimental design [15].

The two modalities with cells and nuclei from stored tissue blocks do not require any cell enrichment steps, though it is possible that either formalin fixation or snap freezing affect cell type proportions differently, as enzymatic and mechanical dissociations processes may also do. Hence, all modalities may have biases that need to be understood. To ameliorate for these biases and to ensure maximal proportional representation, Admati et al. [24] and Garcia-Flores et al. [25] used both cells derived from freshly dissociated samples as well as nuclei from snap frozen samples from the same placentas. It is a valuable strategy, but we note that in other tissues more inherently free from red cell contamination and with highly viable cells directly post dissociation, biases may be less pronounced. Ultimately, the variation in cell type proportions can have severe impact on studies interested in investigating particular cell types, as certain modalities may completely exclude cell types well represented in vivo, such as the absence of cytotrophoblasts (CTB) from the plate based Fixed_in-situ_PolyA, or placental myofibroblasts in the modalities requiring freshly dissociated tissue likely due to the shape of this cell type.

Understanding the levels of variation in the protein coding transcriptomes of the same cell types across modalities allows us to identify which ones produce equivalent results and which, if any, do not. As expected, the Nuclei_droplet_PolyA had the lowest level of correlation with the other droplet-based modalities, most likely due to the lack of cytoplasmic mRNAs inherent in this modality. Surprisingly however, the Fixed_in-situ_PolyA and Nuclei_droplet_PolyA modalities showed the highest level of correlation. This is best observed in the PCA and UMAP presented.

This may suggest that though the Fixed_in-situ_PolyA modality as currently offered by Parse Biosciences and other providers is meant to capture both cytoplasmic and nuclear mRNA, there may be a bias in their chemistry favouring nuclear transcripts. This suggestion is somewhat supported by the observation that the Fixed_in-situ_PolyA had considerably higher levels of expression on non-protein coding nuclear genes (MIR, LINC) than the three whole-cell droplet-based modalities. One potential explanation we have not yet explored is that the specialized fixation and permeabilization required in Fixed_in-situ_PolyA for cells to act as a reaction chamber, through multiple rounds of transcriptomic indexing, also permits cytoplasmic mRNA to escape the cellular boundary. This again would suggest an effect intrinsic to this technology, which may not be resolved either by new versions of chemistry or alternative commercial providers, Scale Biosciences for example. This is an important consideration for studies where cytoplasmic transcripts may be of particular interest, as recently demonstrated in placentas complicated by preeclampsia [15].

Beyond the potential for the resolution of biological samples, single cell transcriptomics is now commonly used in case-control comparison studies to detect disease signatures within and across cell types [2]. We found that modality will impact the proportions of cell types and their comparison. This could render changes in proportions that are not biological but method-dependant and should be taken into consideration in case-control studies. Moreover, the reliability of each method to detect disease markers is of paramount importance. In as much as reliability can be assessed by the reproducibility of DEGs detected across modalities, Fixed_in-situ_PolyA is notably less able to detect DEGs found by other modalities. This effect is increasingly pronounced when exploring the highest and lowest ranked DEGs by fold change. This suggests that even when DEGs are commonly detected across single cell modalities, their ranked position by fold change varies significantly, potentially shifting markers of disease in case-control comparisons. Notably, there is also a pattern of high concordance between Fixed_in-situ_PolyA and Nuclei_droplet_PolyA in the most downregulated genes.

Taken together, we present a comprehensive analysis of the varying effect of different single cell transcriptomic modalities in a challenging tissue, the human placenta, highlighting the strengths and weaknesses inherent in each modality (**Figure 5b**). We provide novel and essential considerations required for experimental design and analytical approaches in single cell transcriptomics, some relevant to the identification of disease markers. These findings are likely applicable to other challenging tissues, whether due to inherent biological characteristics, sampling approaches available or degrading effects of pathology.

## ACKNOWLEDGEMENTS

We thank Sasha Tunnock, Tyrrell Hatch and Claire Boevink, research midwives at University College London Hospitals, NHS Foundation Trust. Also, Dr Diego Chillon Pino for scientific input and the MRC Laboratory of Medical Sciences (LMS) Genomics Facility for the processing of the *in situ* spatial transcriptomic slide. Finally we would like to thank UCL Genomics for providing the sequencing of single cell libraries.

## AUTHOR CONTRIBUTIONS

TX conceived of the project, designed the and performed the experiments and performed data interpretation and writing of the manuscript. JJMV and GH provided advice and reviewed the manuscript. SLH recruited the patients, assisted in tissue sampling, provided advice and reviewed the manuscript. YESC provided advice and reviewed the manuscript and assisted in supervision. SC conceived of and supervised the project, contributed to data interpretation and writing of the manuscript.

## CONFLICT OF INTEREST

There are no competing interests to declare.

## FUNDING

This work was supported by the MRC (MR/W028158) and the NIHR GOSH Biomedical Research Center (NIHR203326).

## Notes

### Competing Interest Statement

The authors have declared no competing interest.

