## Supplementary figures for "The challenges of single cell transcriptomics on difficult human tissue: the placenta"

a)

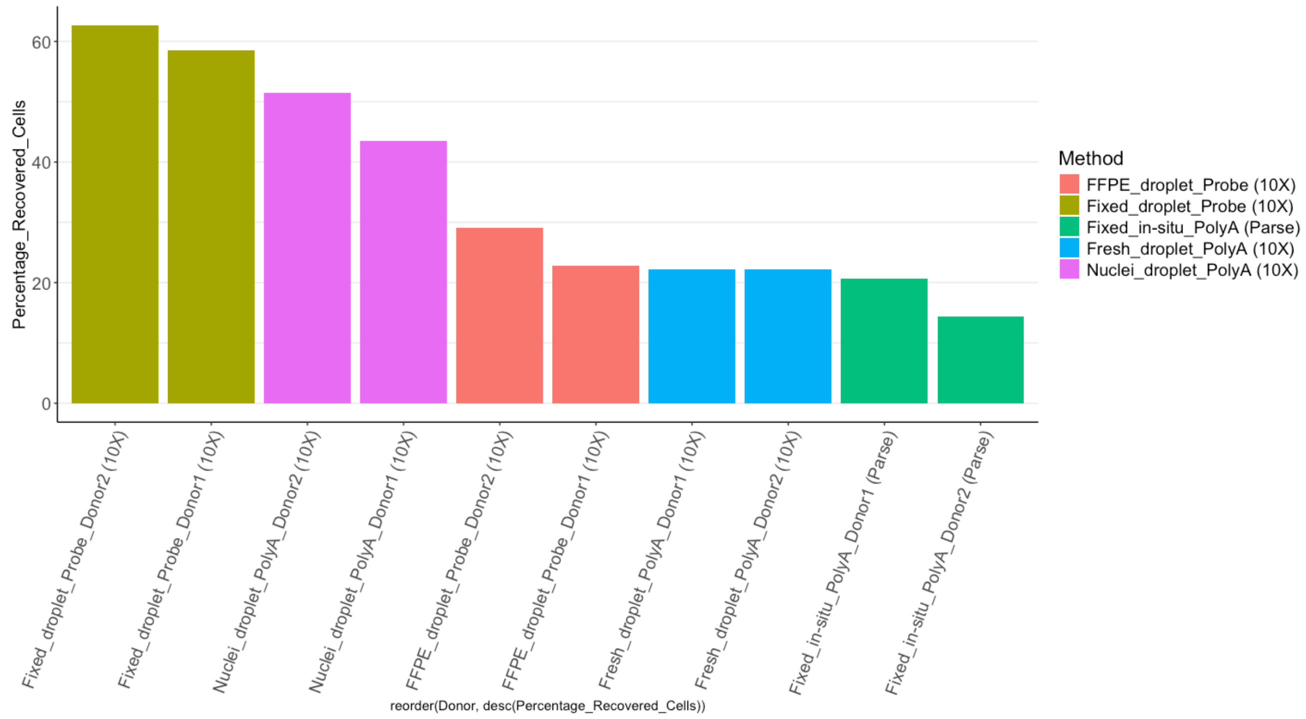


b)

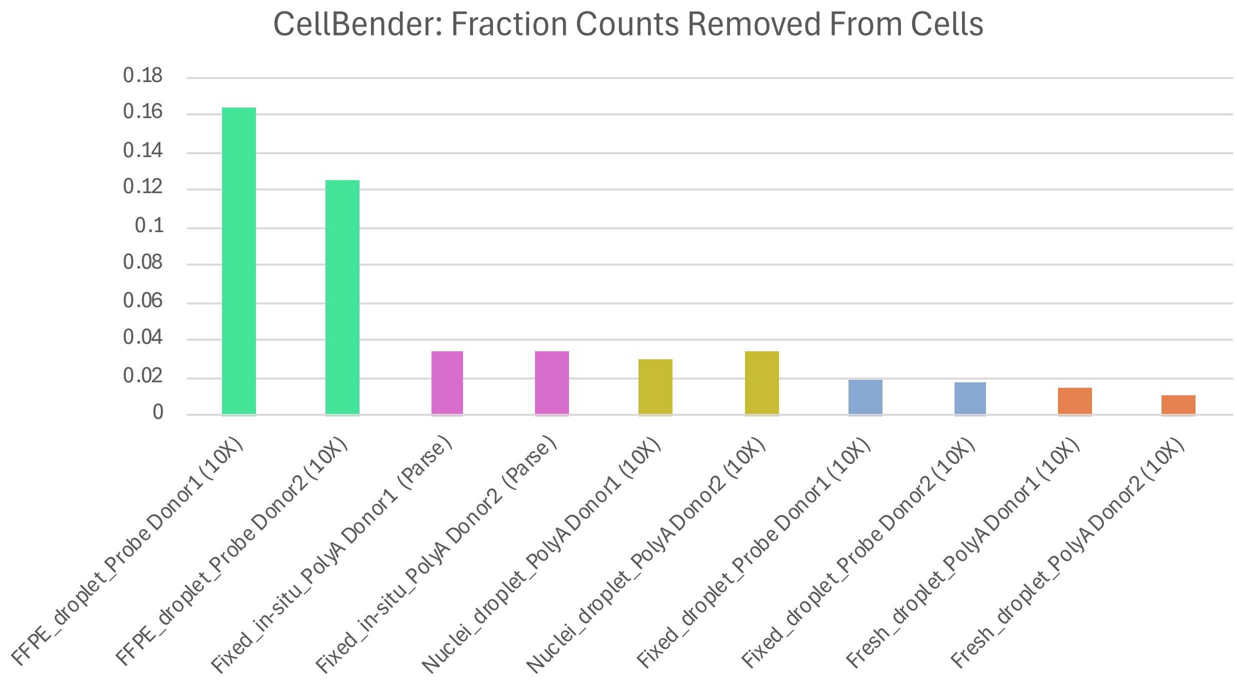


**Extended figure 1.** Performance metrics per biological replicate. **a**) Capture efficiency, showing the percentage of cells used for analysis post QC filtering in relation to cells inputted for each biological replicate. **b**) Background, illustrating per biological replicate the fraction of reads removed by the computational tool used for removing reads assigned to ambient RNA.

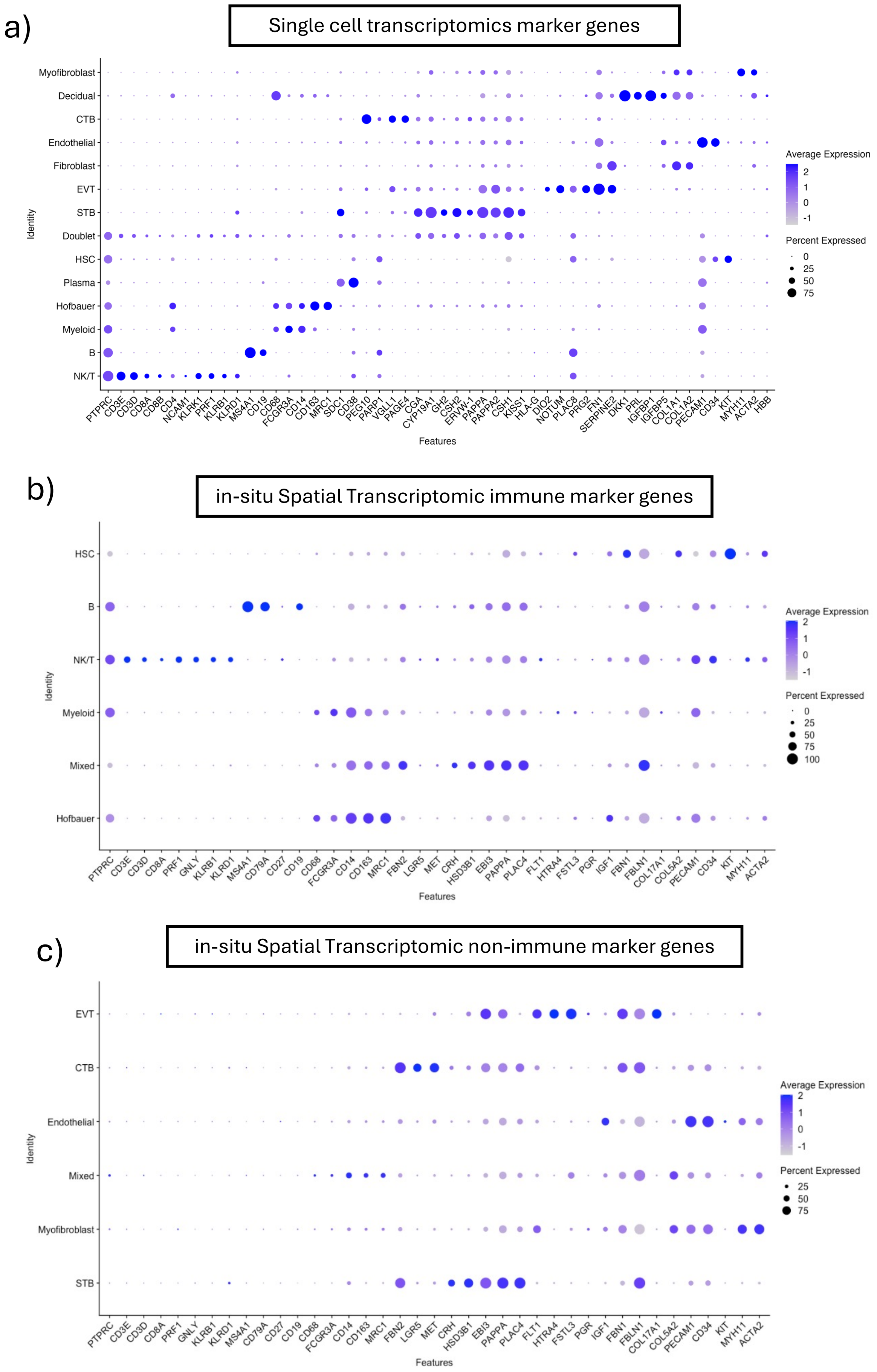


**Extended figure 2.** Marker gene expression of annotated cells. **a**) Dotplot illustrating marker gene expression of annotated cell types in combined dataset including all five modalities. **b**) Dotplot illustrating marker gene expression of annotated cell types in immune compartment of in-situ Spatial Transcriptomic sample. **c**) Dotplot illustrating marker gene expression of annotated cell types in non-immune compartment of in-situ Spatial Transcriptomic sample.

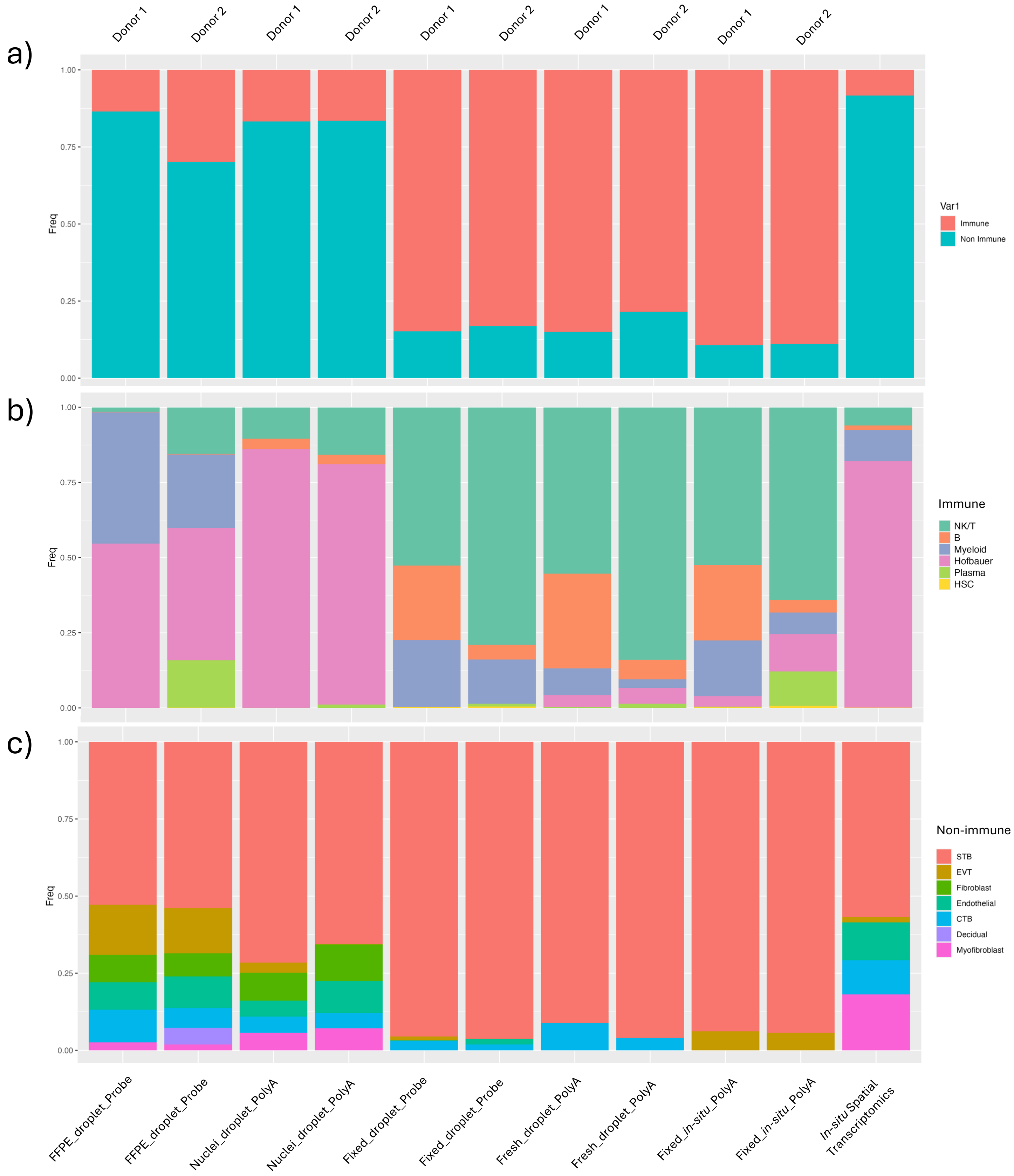


**Extended figure 3.** Cell type proportions by biological replicate. **a**) Relative proportion of cells annotated as immune and non-immune cells by biological replicate and modality. Relative proportions of cell annotated within the **b**) immune and **c**) non-immune cell types by biological replicate and modality.

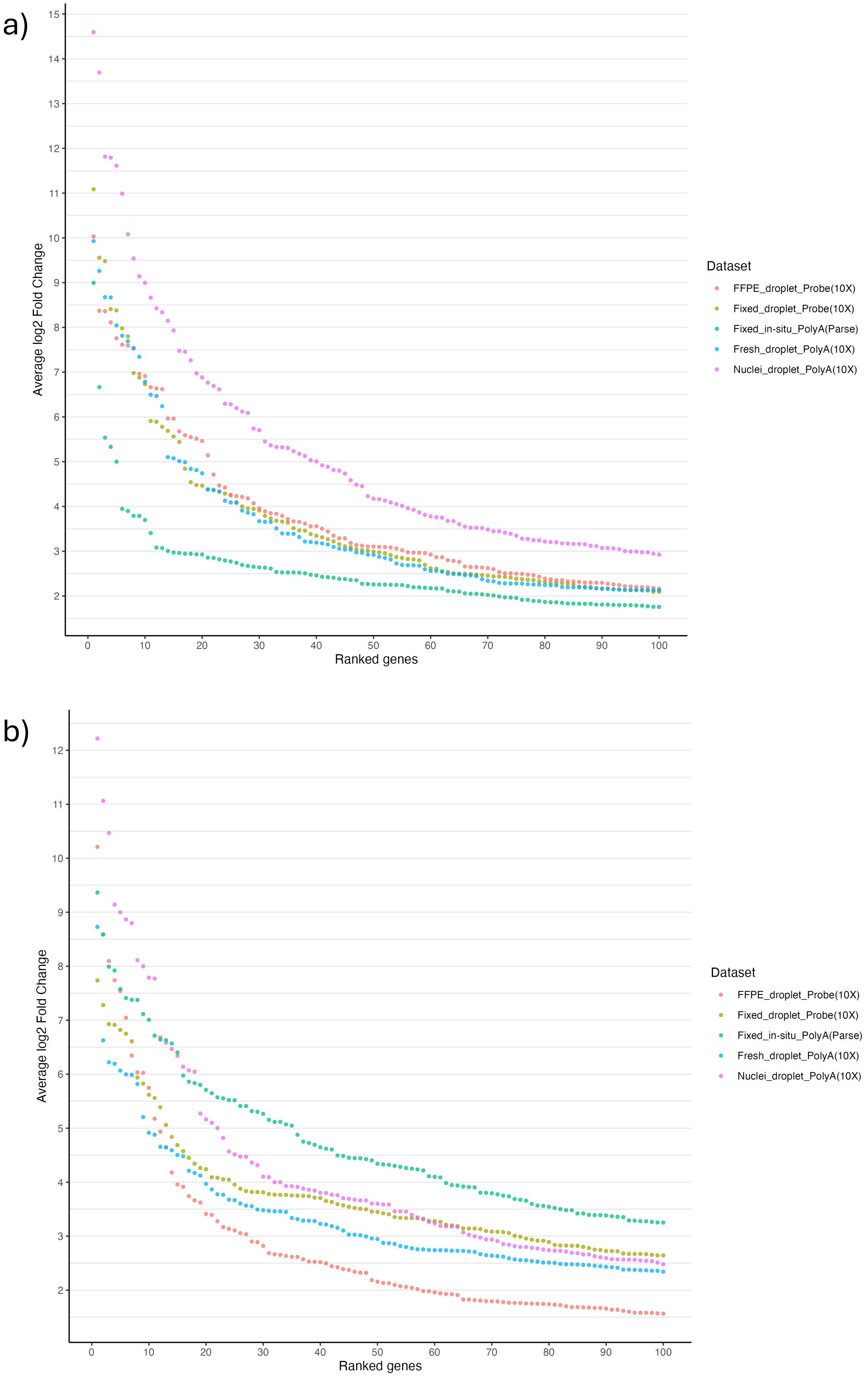


**Extended figure 4.** Ranking of differentially expressed genes per modality. **a**) Top 100 over expressed genes ranked by average log2 fold change (adjusted p-value <0.05), when comparing Syncitiotrophoblasts (STBs) between biological replicates, within each modality. **b**) Top 100 under expressed genes ranked by average -log2 fold change (adjusted p-value <0.05), when comparing Syncitiotrophoblasts (STBs) between biological replicates, within each modality.

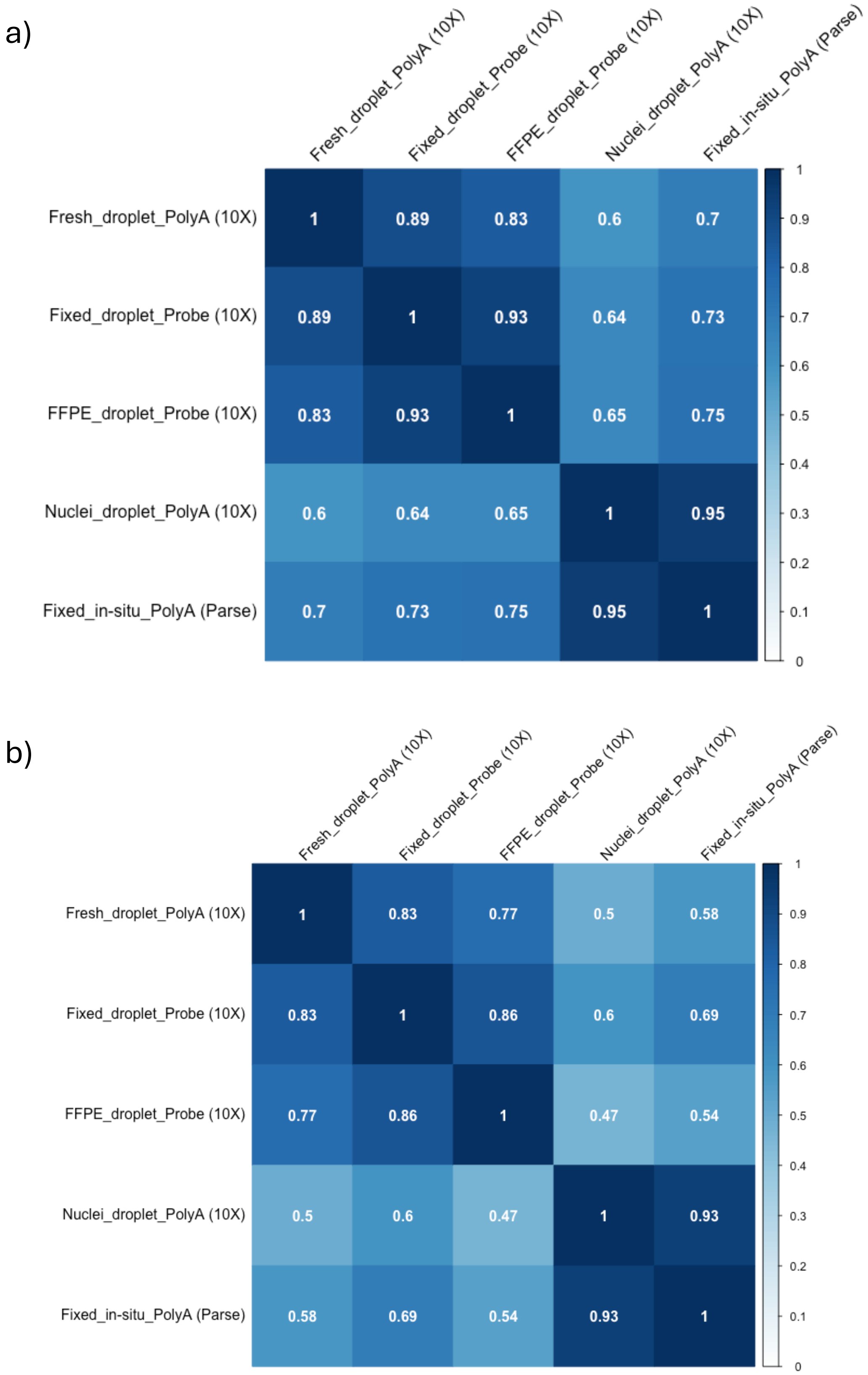


**Extended figure 5.** Correlation matrix of pseudobulked protein coding gene expression of **a**) STBs and **b**) Natural Killer/T-cells across modalities.
